# Relative Patlak Plot for Dynamic PET Parametric Imaging Without the Need for Early-time Input Function

**DOI:** 10.1101/268730

**Authors:** Yang Zuo, Jinyi Qi, Guobao Wang

## Abstract

The Patlak graphical method is widely used in parametric imaging for modeling irreversible radiotracer kinetics in dynamic PET. The net influx rate of radiotracer can be determined from the slope of the Patlak plot. The implementation of the standard Patlak method requires the knowledge of full-time input function from the injection time until the scan end time, which presents a challenge for use in the clinic. This paper proposes a new relative Patlak plot method that does not require early-time input function and therefore can be more efficient for parametric imaging. Theoretical analysis proves that the effect of early-time input function is a constant scaling factor on the Patlak slope estimation. Thus, the parametric image of the slope of the relative Patlak plot is related to the parametric image of standard Patlak slope by a global scaling factor. This theoretical finding has been further demonstrated by computer simulation and real patient data. The study indicates that parametric imaging of the relative Patlak slope can be used as a substitute of parametric imaging of standard Patlak slope for certain clinical tasks such as lesion detection and tumor volume segmentation.

## 1. Introduction

Dynamic positron emission tomography (PET) provides four-dimensional distribution (threedimensional space plus one-dimentional time) of radiotracer in living body and is attracting more and more research interests [Schmidt and Turkheimer, 2002, Rahmim et al, 2009, Wang and Qi, 2013, Reader and Verhaeghe, 2014]. Analyzing dynamic PET data relies on kinetic modeling which commonly uses a temporal model to describe the kinetics of radiotracer uptake [Schmidt and Turkheimer, 2002]. Voxel-wise implementationof kinetic modeling provides parametric maps of kinetic parameters indicating the biological characters of the tissue [Gunn et al., 1998]. Parametric imaging has been found useful in many applications including tumor detection [Kordower et al., 2000, Gill et al., 2003] and extraction of metabolic tumor volume (MTV) [Visser et al., 2008].

The Patlak graphical plot is a widely used kinetic analysis method in dynamic PET for extracting the net influx rate of irreversible uptake of a radiotracer [Patlak et al., 1983, Patlak and Blasberg, 1985]. It is also used for kinetic modeling in dynamic magnetic resonance imaging (MRI) [Hackstein et al., 2003] and computed tomography (CT) [Hackstein et al., 2004, Miles et al., 1999, Hom et al., 2009]. Compared with nonlinear compartmental modeling, the Patlak method uses a linear model and has the advantage of being computationally efficient for parametric imaging and being easier to be implemented into new reconstruction methods [Wang et al., 2008, Tsoumpas et al., 2008, Tang et al., 2010, Angelis et al., 2011, Zhu et al., 2013] and for whole-body imaging [Karakatsanis et al., 2015, Karakatsanis et al., 2016, Zhu et al., 2013, Hu et al., 2017].

Input function is essential for tracer kinetic modeling. Although the Patlak graphical plot only examines the time points of tissue activity at steady state, the standard Patlak method requires the knowledge of full-time input function from the radiotracer injection time until the dynamic scan end time. To obtain the information of full-time input function, either many blood samples are needed if arterial blood sampling is used or a long scan covering early-time points is required if the image-derived or reference region input function is used. For example in dynamic ^18^F-FDG PET imaging, an one-hour dynamic scan is required to derive the full blood input function from dynamic images, though only late 20-30 minutes are actually used to extract the tracer activity of tissue. This requirement for early-time input function presents a challenge for applying the Patlak method in the clinic, particularly for whole-body imaging [Hu et al., 2017].

This paper proposes a new relative Patlak plot method for PET parametric imaging. Compared with the standard Patlak plot, the proposed relative plot does not require the information of early-time input function. Mathematical analysis is used to show the relationship between the slope of the new plot and that of the standard Patlak plot. The theoretical findings are further validated by computer simulation of dynamic PET data and real patient scan data.

## 2. Theory

### 2.1. Standard Patlak Plot

Let us denote the radiotracer concentration at time *t* in a tissue region of interest (ROI) or voxel by *C_T_*(*t*) and tracer concentration in the plasma by *C_P_*(*t*). The Patlak plot exploits the linearity between the normalized tissue concentration and normalized integral of input function *C_P_*(*t*) after a steady-state time *t*\*. Mathematically, it is described by a linear equation:

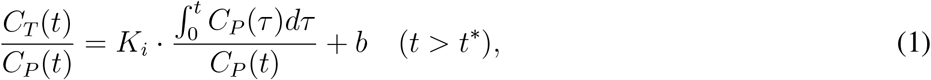

where *K*_*i*_ is the slope constant that represents the net influx rate of irreversible uptake of a radiotracer and *b* is the intercept which equivalently indicates blood volume in the tissue and the normalized tracer concentration from reversible compartments.

By acquiring dynamic PET data for multiple time frames, one can plot the data of

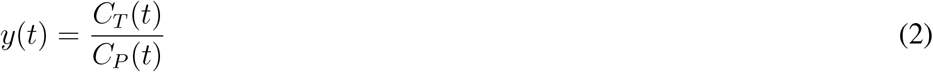

and

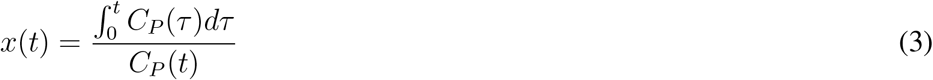

of each time frame and fit the data points using the linear model Eq. (1). The slope *K*_*i*_ and intercept *b* are then estimated by the linear regression. Note that although *y*(*t*) is only sampled for *t* > *t*\*, *x*(*t*) contains the integral of the input function *C_P_*(*t*) from the injection time *t* = 0 till the scan end time. Thus the full-time input function needs to be known for the standard Patlak plot.

Given a set of measurements at *M* time points 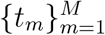 with *t*_1_ = *t*\*, the Patlak slope and intercept can be estimated using the following least-squares formulation,

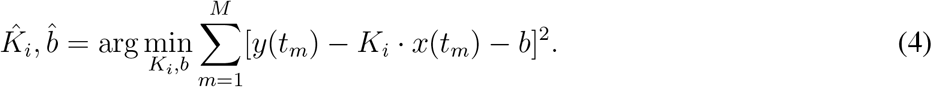

The optimal solution has the following analytic formula

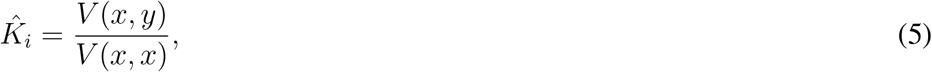

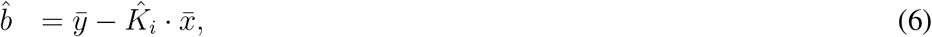

where *V*(·, ·) and 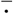 have the forms as follows

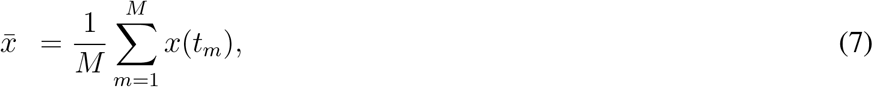

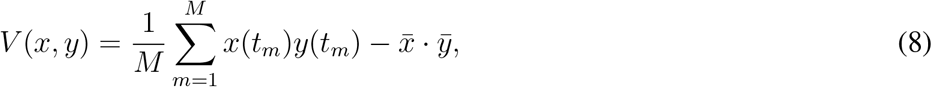

which correspond to the mean and covariance if *x* and *y* are considered as random variables.

### 2.2. Proposed Relative Patlak Plot

We propose a new relative Patlak plot which has the following model equation:

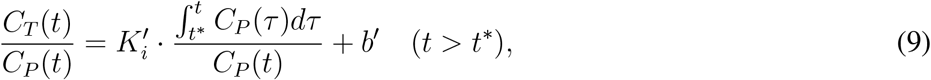

where 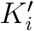 and *b′* are the slope and intercept of the new plot, respectively.

The new relative Patlak method plots the data of *y*(*t*) in Eq. (2) versus

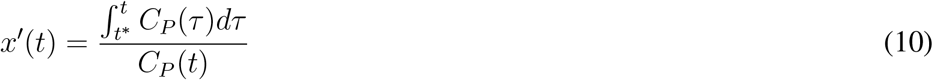

to get the slope 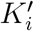 and intercept *b′*. This new model is very similar to the standard Patlak plot equation, except that the integral of the input function *C_P_*(*t*) here is from *t*\* to *t*, not from 0 to *t*. The integral of *C_P_*(*t*) over early time from 0 to *t*\* is no longer needed in this new plot.

Similarly to the least-squares estimation for the standard Patlak plot, the slope and intercept of the relative Patlak plot can be estimated by

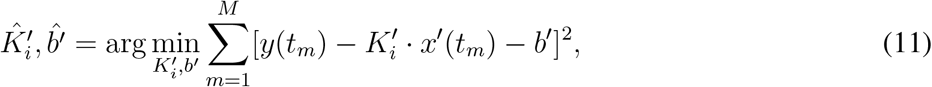

which gives the following optimal solution

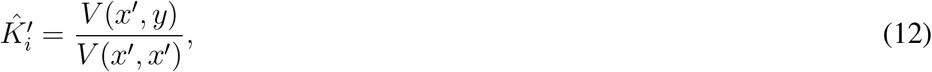

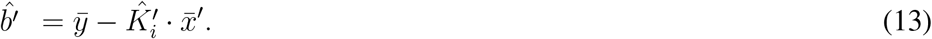

where 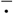 and *V*(·, ·) are defined in Eqs. (7) and (8), respectively.

### 2.3. Theoretical Relation Between the Two Plots

The new relative Patlak plot is closely related to the standard Patlak plot. Here we examine the theoretical relation between the two plots using analytical derivations. Let us define

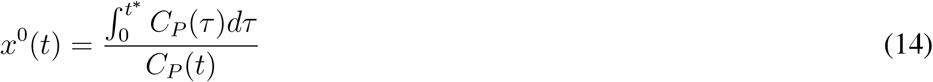

to account for the component of early-time input function which appears in *x*(*t*) but not in *x′*(*t*). Obviously, we have

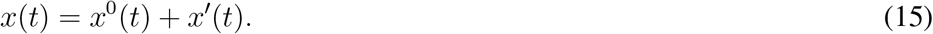

Without loss of generality in dynamic PET, the late-time input function *C_P_*(*t*) for *t* > *t*\* can be analytically expressed by an exponential function:

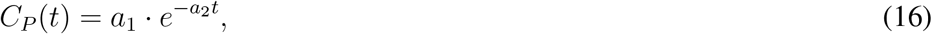

where *a*_1_, *a*_2_ > 0. For example, the widely used Feng model [Feng et al., 1993] has the form

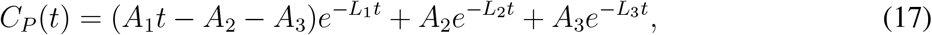

with the following parameters *A*_1_ = 851.1 mg/100mL/min, *A*_2_ = 20.8 mg/100mL, *A*_3_ = 21.9 mg/100mL, *L*_1_ = 4.1 min^−1^, *L*_2_ = 0.01 min^−1^, *L*_3_ = 0.12 min^−1^. For *t* > *t*\* = 30 minutes, the Feng input function is dominated by the term 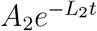 while the other two terms are negligible. As a result, *x*^0^(*t*) and *x′* (*t*) can be rewritten as

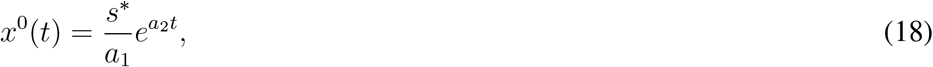

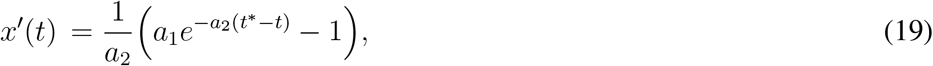

with *s*\* being the integral of blood input over the early time from time 0 to *t*\*,

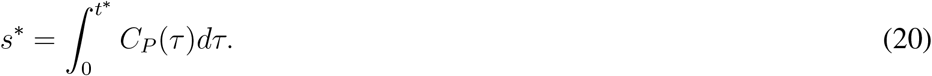

It is then not difficult to verify the following linear relationship between *x*^0^(*t*) and *x′*(*t*):

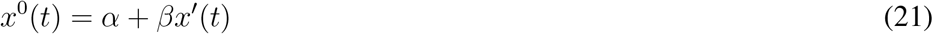

where *α* and *β* are both constants that only depend on *t*\*:

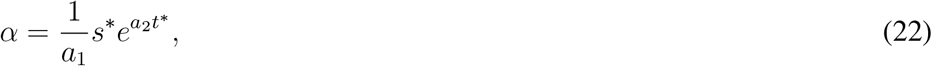

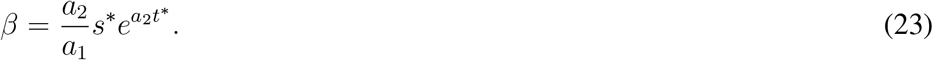

Using Eq. (15), the standard Patlak plot model Eq. (1) can be re-written as

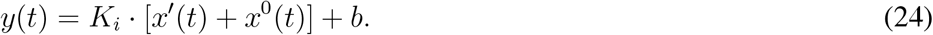

Substituting Eq. (21) into Eq. (24), we obtain the following equivalence,

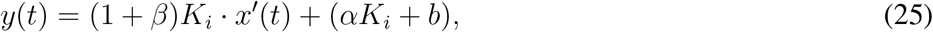

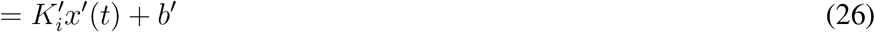

with

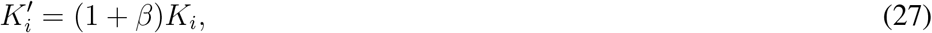

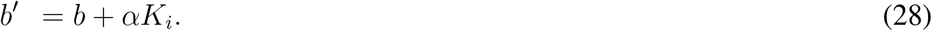

The two equations above indicate that the relative Patlak slope 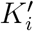 is proportional to the standard Patlak slope *K*_*i*_ with a scaling factor (1 + *β*). The intercept of the relative Patlak plot is equivalent to the intercept of the standard Patlak plot plus a shift α*K*_*i*_.

The scaling factor (1 + *β*) only depends on the input function and is independent of tissue time activity. It is therefore a global scaling factor when the Patlak plot is implemented for parametric imaging. Thus, the parametric image of the relative Patlak slope 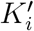 is equivalent to the parametric image of the standard Patlak slope *K*_*i*_ up to a scaling factor.

### 2.4. Theoretical Relation Between the Least Squares Estimates

In practice, the slope and intercept of a graphical plot are commonly estimated by a least squares optimization. Here we examine the theoretical relation between the standard Patlak and relative Patlak least squares estimates.

Based on Eq. (21), *x*(*t*) and *x′*(*t*) approximately satisfy a linear relation:

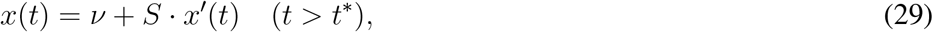

where *S* and *v* are respectively equivalent to (1 + *β*) and *α* if the late-time blood input is described by an exponential function. Alternatively, *S* and *v* can be estimated by a linear regression without assuming an exponential model:

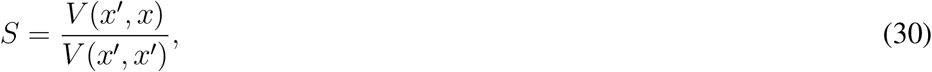

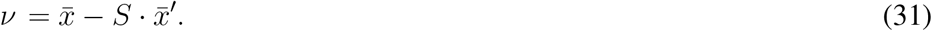

Substituting Eq. (29) into the least squares estimate of the standard Patlak slope 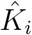 in Eq. (5) leads to the following expression,

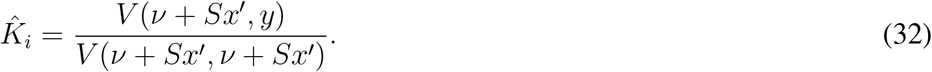

Note that the function *V*(·, ·)has the following properties:

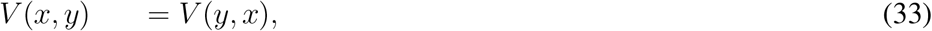

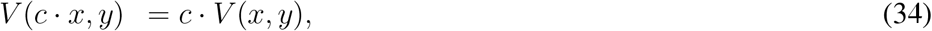

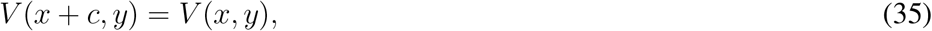

where *c* is an arbitrary constant. Using these properties and the definition of the least-squares estimate of the relative Patlak slope 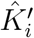 defined in Eq. (12), we then obtain the scaling relation between 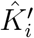 and 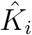 :

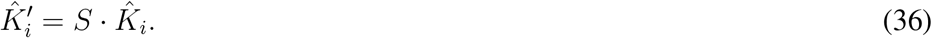

We can also derive the following relation between the least-squares estimates of the two intercepts 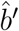 and 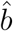:

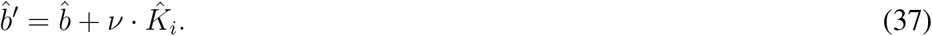

These results indicate that the theoretical relation between the standard Patlak plot and relative Patlak plot holds true for their least squares estimation.

## 3. Materials and Methods

### 3.1. Validation Using Computer Simulation

We first conducted a computer simulation to validate the theoretical results on the scaling relationship between the standard Patlak slope and relative Patlak slope. One-hour dynamic ^18^F-FDG scan was simulated following the scanning sequence of a total of 55 frames: 30 × 10- second frames, 10 × 60-second frames and 15 × 180 -second frames. The blood input function in this simulation was generated using the analytical Feng model [Feng et al., 1993]. Following the standard two-tissue compartmental model, we simulated 10, 000 groups of random kinetic parameters which follow a Gaussian distribution with the mean of kinetic parameters being *K*_1_ = 0.81 mL/mL/min, *k*_2_ = 0.38 min^−1^, *k*_3_ = 0.1 min^−1^, *k*_4_ = 0 min^−1^ and standard deviation being 40% of the mean kinetic parameters. Noise-free time activity curves (TACs) were generated with the simulated FDG kinetics and blood input function. Zero-mean Gaussian noise were then added to each noise-free TAC 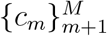 using the noise standard deviation [Wu and Carson, 2002] defined by

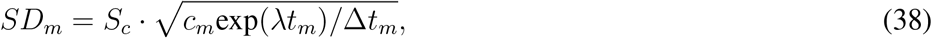

where *S*_*c*_ is a scale factor to adjust *SD* to match with realistic dynamic FDG-PET data at different noise levels. *S*_*c*_ = 1.0 was used to simulate a voxel-level high noise in this simulation. λ is the decay constant of the radiotracer set to be ln(2)/*T*_1/2_ with *T*_1/2_ = 109.8 minutes. Δ*t*_*m*_ is the scan duration of time frame *m* and *t*_*m*_ is the middle time of frame *m*.

### 3.2. Validation Using Patient Scans

We further validated the theoretical results using dynamic FDG-PET scans of two human patients, one with breast cancer and the other with coronary heart disease.

The breast patient scan was operated on the GE Discovery 690 PET/CT scanner at UC Davis Medical Center. The patient received 5 mCi ^18^F-FDG with a bolus injection. Listmode time-of-flight data acquisition commenced right after the FDG injection and lasted for 60 minutes. A low-dose transmission CT scan was then performed at the end of PET scan to provide CT image for PET attenuation correction. The raw data were then binned into a total of 49 dynamic frames: 30 × 10 seconds, 10 × 60 seconds and 9 × 300 seconds. Dynamic PET images were reconstructed using the standard ordered subsets expectation maximization (OSEM) algorithm with 2 iterations and 32 subsets as provided in the vendor software. All data corrections including normalization, attenuation correction, scattered correction and randoms correction, were included in the reconstruction process. A region of interest was placed in the left ventricle region to extract blood input function.

**Figure 1.**
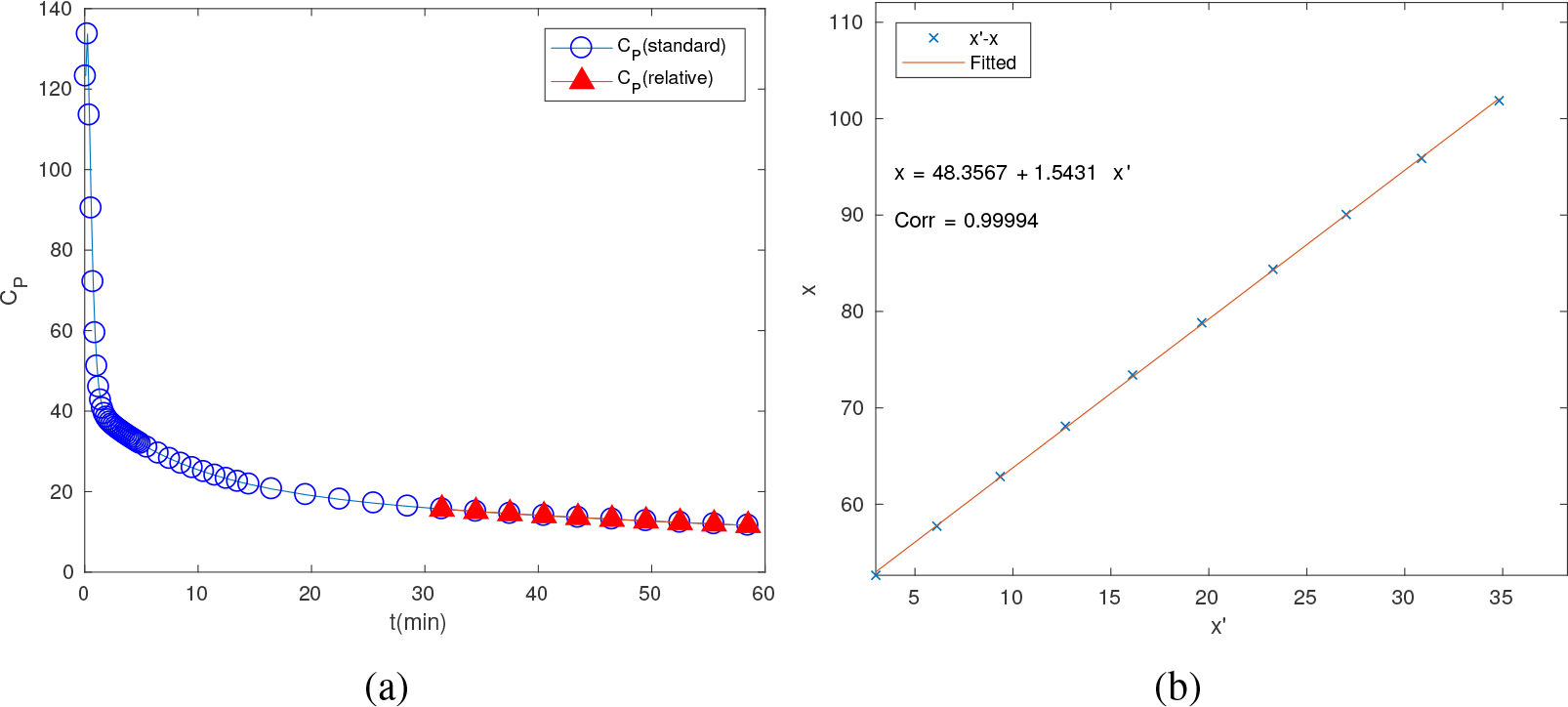
Blood input functions in the simulation. (a) Full-time blood input function (circles) for the Standard Patlak and late time points (solid triangles) for the relative Patlak; (b) Linear relation between *x* and *x′* with *t*\* = 30 minutes.

The cardiac scan was performed on the GE Discovery ST PET/CT scanner at UC Davis Medical Center in two-dimensional mode. The scanner has no time-of-flight capability. The patient received 10 mCi ^18^F-FDG with a bolus injection. List-mode data acquisition commenced right after the FDG injection and lasted for 60 minutes. A low-dose transmission CT scan was then performed at the end of PET scan to provide CT image for PET attenuation correction. The raw data were binned into a total of 49 dynamic frames: 30 × 10 seconds, 10 × 60 seconds and 9 × 300 seconds. Dynamic PET images were reconstructed using the standard ordered subsets expectation maximization (OSEM) algorithm with 2 iterations and 30 subsets as provided in the vendor software. All data corrections including normalization, attenuation correction, scattered correction and randoms correction, were included in the reconstruction process. The blood input function was also extracted from the left ventricle region.

## 4. Results

### 4.1. Simulation Results

The simulated noisy TACs were first analyzed using the standard Patlak plot and new relative Patlak plot with a start time *t*\* = 30 minutes. For the standard Patlak plot, the full-time blood input from 0 to 60 minutes was used to estimate the slope *K*_*i*_ For the relative Patlak plot, only the input function after *t*\* was used to estimate the slope 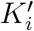. Early-time input function is not needed for the relative Patlak plot, which is equivalent to setting those time points to zeros. The two input functions are graphically compared in Fig. 1(a). We then verified the linear relation between 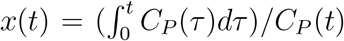 and 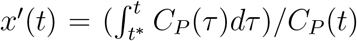 with *t* > *t*\*. Figure 1(b) shows the plot of *x*(*t*) versus *x′*(*t*) for the simulated Feng input function. The linear fit was excellent with a pairwise linear correlation coefficient close to 1.

**Figure 2.**
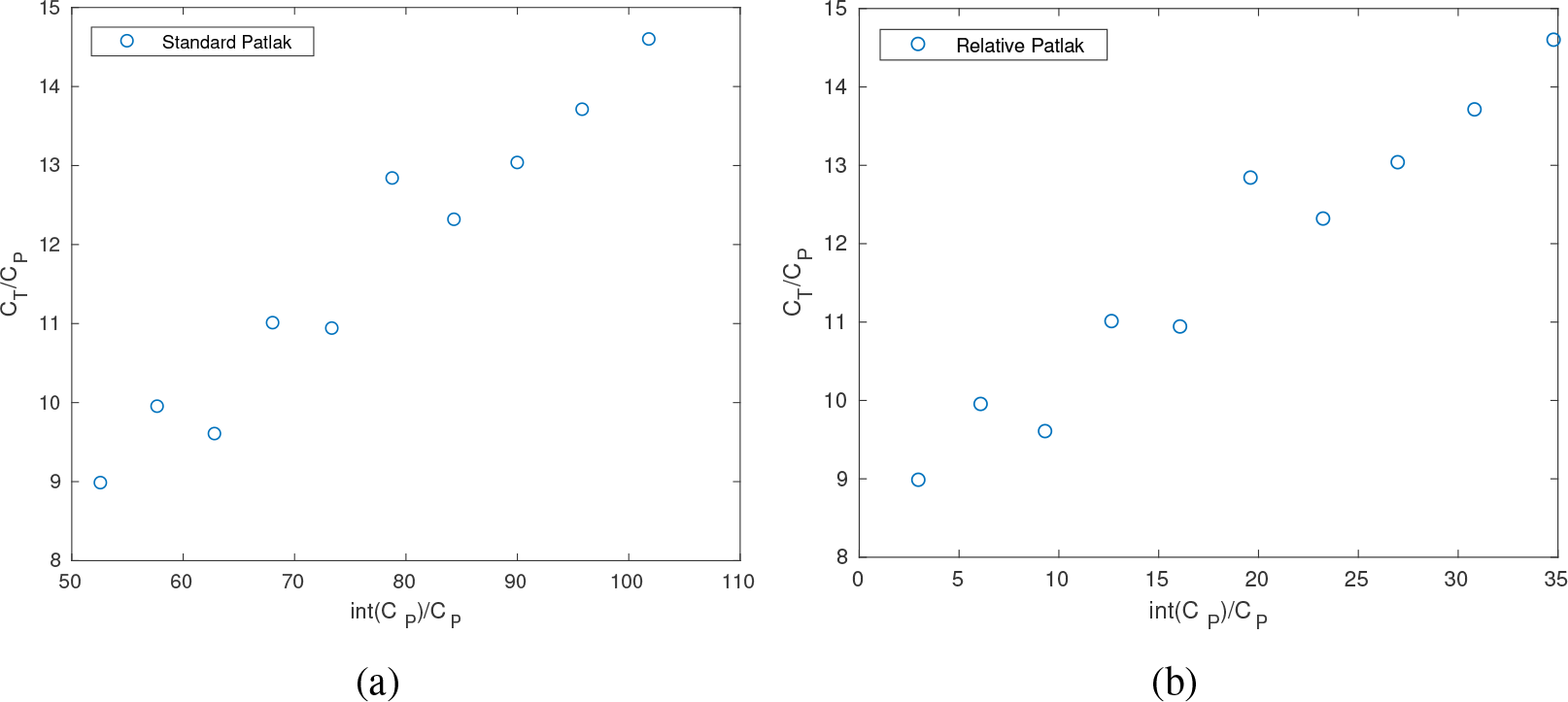
Comparison of the standard Patlak plot and relative Patlak plot in the simulation. (a) standard Patlak plot; (b) relative Patlak plot.

**Figure 3.**
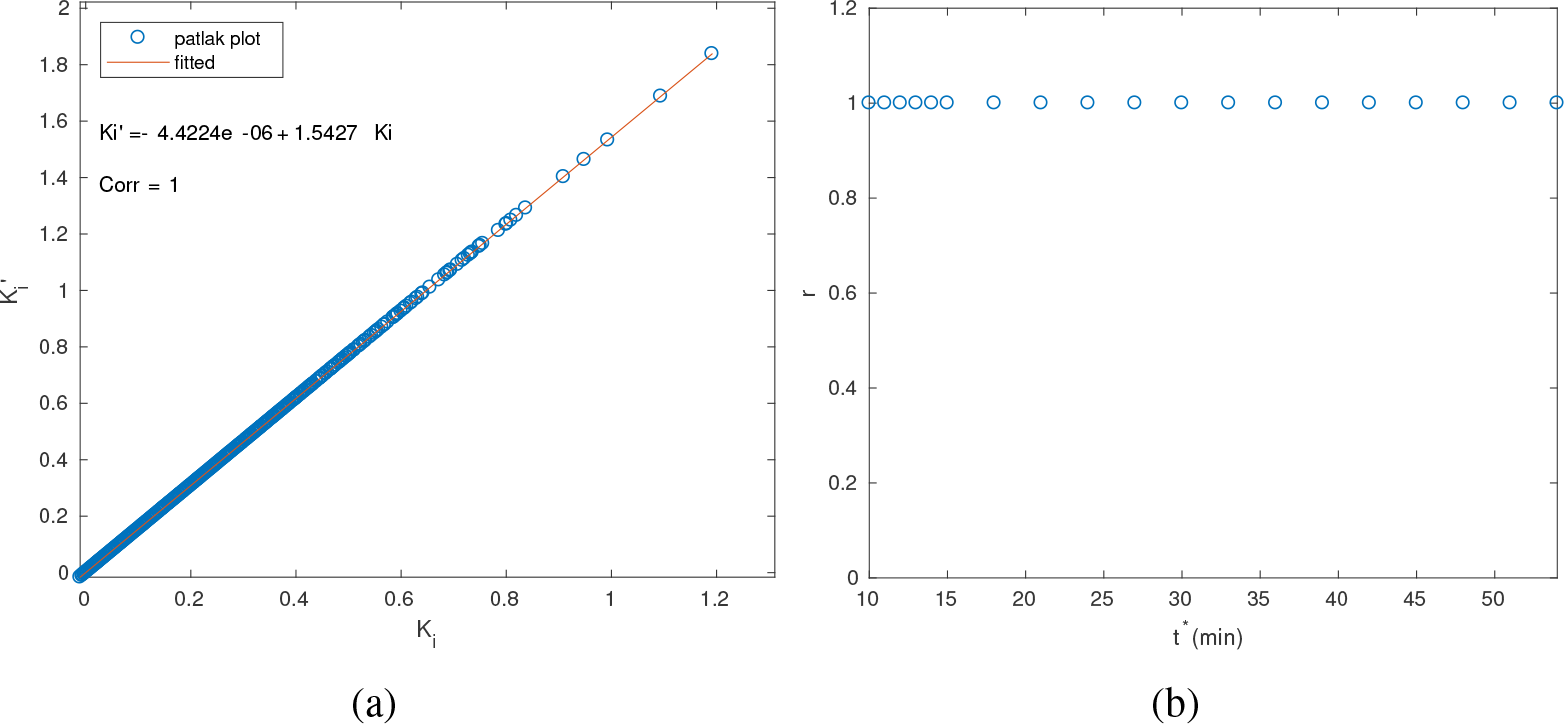
Results of simulation. (a) Relation between 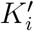 and *K*_*i*_; (b) Correlation coefficient of 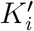 versus *K*_*i*_, for various *t*\* values.

Examples of the the standard Patlak plot and relative Patlak plot are shown in Fig. 2 (a) and (b). We examined the linearity between the standard Patlak slope *K*_*i*_ and the relative Patlak slope 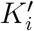. Fig. 3(a) shows the estimated 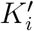 versus *K*_*i*_ values for all the simulated 10, 000 TACs. The correlation coefficient between 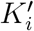 and *K*_*i*_ was 1.0, indicating a perfect linearity. The intercept is negligible, indicating 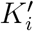 values are equal to *K*_*i*_ values times the scaling factor. The slope of the linear plot of 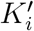 versus *K*_*i*_ is approximately equal to the slope of the linear plot of *x*(*t*) versus *x′*(*t*). The relation between 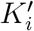 and *K*_*i*_ is therefore verified by the simulation data. Fig. 3(b) further shows that the correlation coefficient between 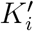 and
*K*_*i*_ remains stable and close to 1 when *t\** varied from 10 minutes to 54 minutes, though the scaling factor between them depends on *t\**. Note that *t\** could not be greater than 54 minutes given the defined time frames, otherwise less than two time points could be used for the Patlak plots.

**Figure 4.**
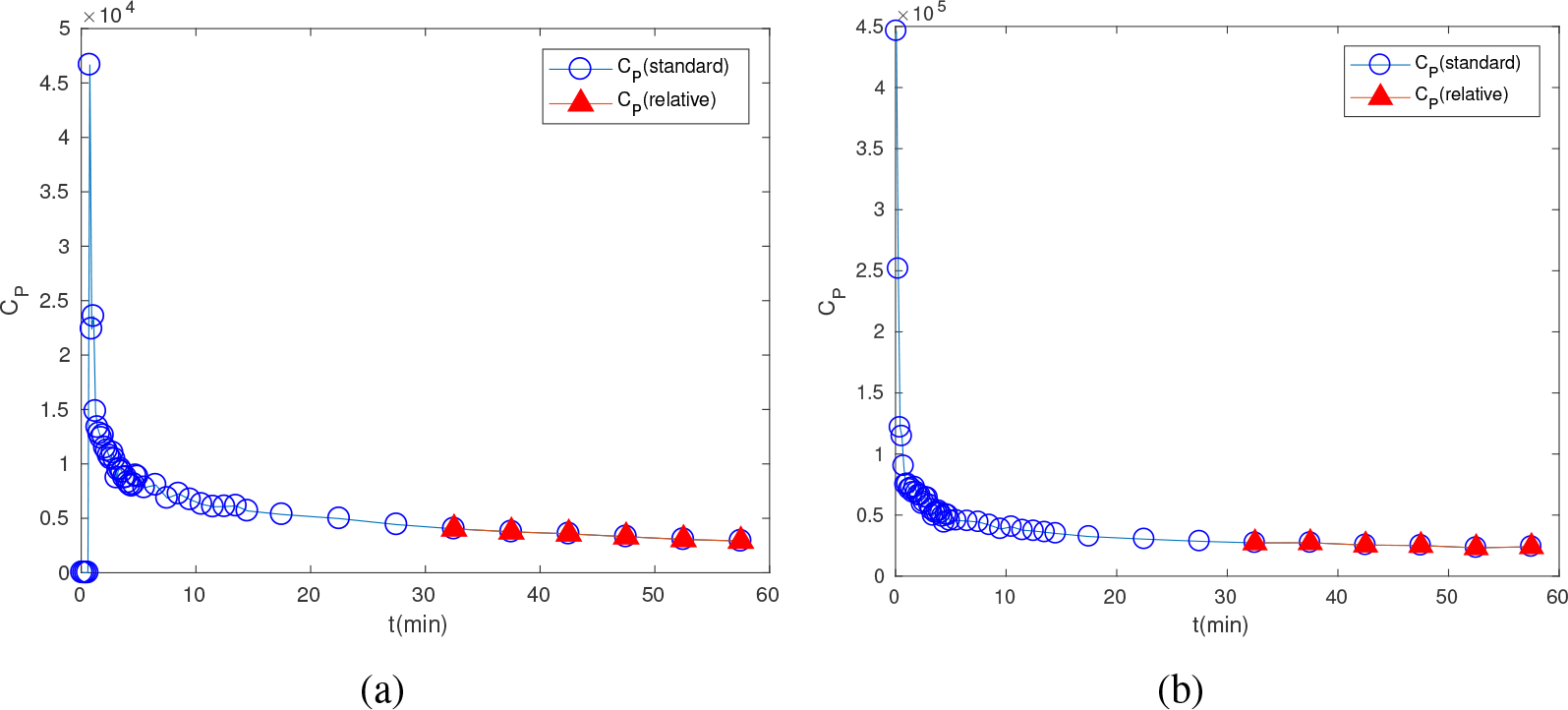
Blood input functions from dynamic FDG-PET scans of human patients. (a) breast cancer patient; (b) cardiac patient.

**Figure 5.**
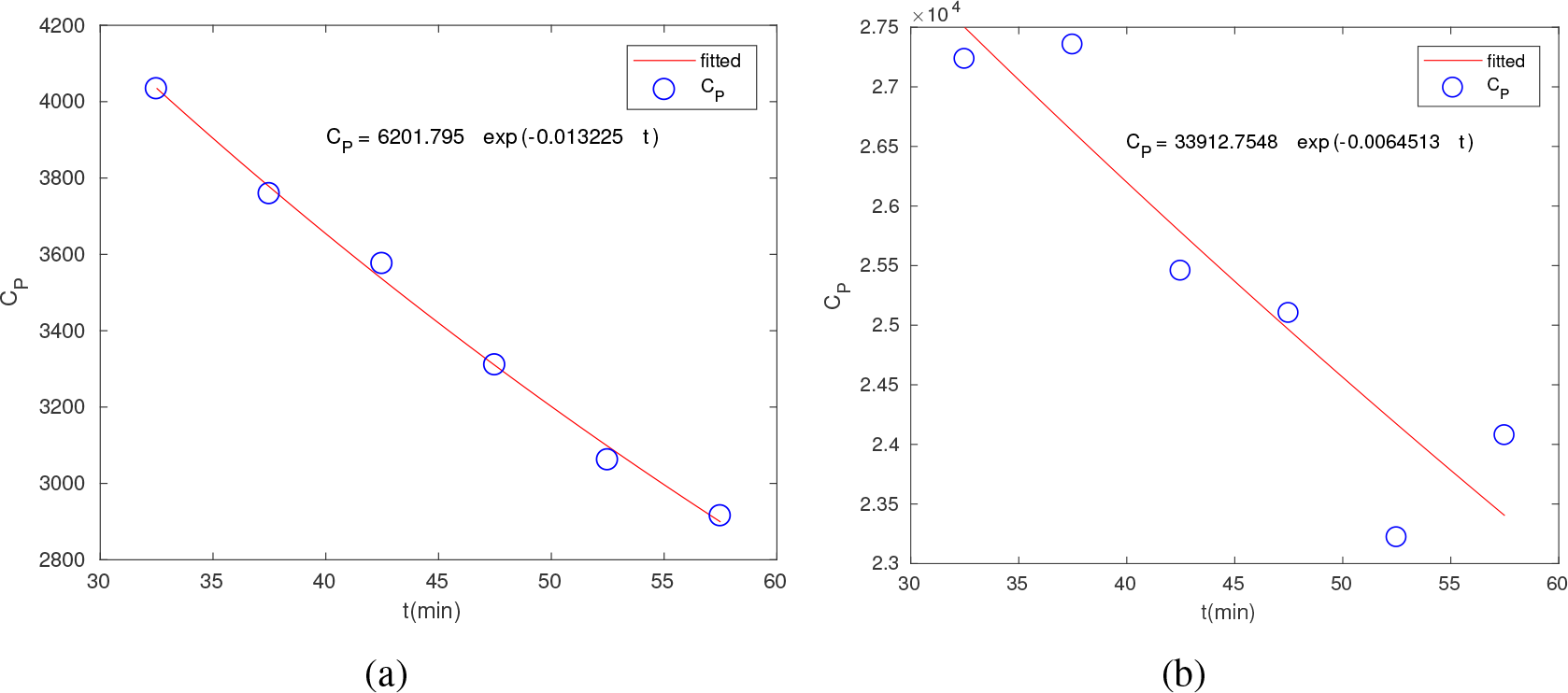
Validation of late-time points of real patient blood input functions fitted by an exponential function. (a) breast patient data, (b) cardiac patient data.

### 4.2. Patient Results

#### 4.2.1. Blood Input Functions

The image-derived input functions from the breast patient scan and cardiac patient scan are shown in Fig. 4(a) and Fig. 4(b), respectively. The start time *t\** was initially set to 30 minutes. Fig. 5 validates that the late-time time points of the two blood input functions can be fitted with the one-exponential function 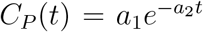 for *t* > *t\**. It is not surprising that higher noise presents in the cardiac patient data because the scan was operated in a 2D mode and also without time-of-flight capability.

**Figure 6.**
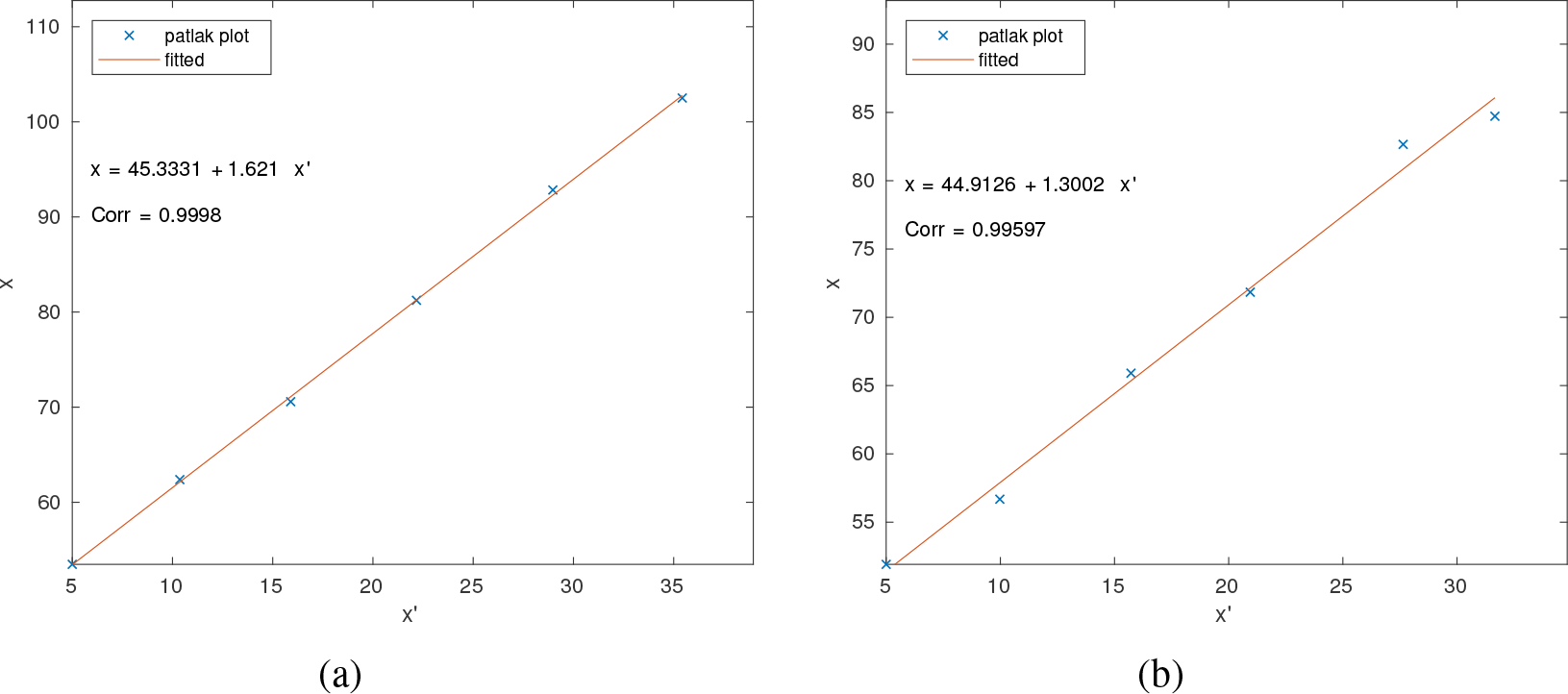
Linear relation between *x*(*t*) and *x′*(*t*) for patient data with *t* ≥ *t*\* = 30 minutes. (a) breast patient data, (b) cardiac patient data.

The linear relation between *x*(*t*) and *x′*(*t*) after *t\** = 30 minutes is shown in Fig. 6(a) for the breast patient data and in Fig. 6(b) for the cardiac patient data.

#### 4.2.2. Breast Patient Result

The parametric maps of *K*_i_ by the standard Patlak model and 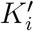 by the relative Patlak model are shown in Fig. 7 for transverse, sagittal and coronal planes. These two parametric images have different absolute values but they appear to be proportional to each other. The plot of 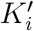 versus *K*_i_ is shown in Fig. 8(a). It is clear that 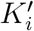 was linearly related to *K*_i_ with a slope of 1.6199 and intercept of 1.7345 × 10^−7^. The intercept was negligible so the linear relation was simply a scaling. The slope of the linear plot of 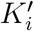 versus *K*_i_ is very close to the slope of the linear plot of *x*(*t*) versus *x′*(*t*). The correlation coefficients between 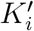 and *K*_i_ was close to 1. Fig. 8(b) further shows the correlation coefficient between 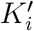 and *K*_*i*_ versus various *t\** values ranging from 10 minutes to 50 minutes. High correlation remains between 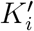 and *K*_i_.

#### 4.2.3. Cardiac Patient Result

The parametric maps of *K*_*i*_ by the standard Patlak model and 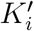 by the relative Patlak model with *t\** = 30 minutes are shown in Fig. 9. Again, the two parametric images appear to be proportional to each other, though with different absolute values. The plot of 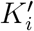 versus *K*_*i*_ is shown in Fig. 10(a). It is again clear that 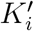 was linearly related to *K*_*i*_ with a slope of 1.2898 and intercept of 7.7 × 10^−6^. The negligible intercept indicates that the linear relation was a scaling. The slope of plotting 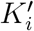 versus *K*_*i*_ was very close to the slope 1.3022 of plotting *x*(*t*) versus *x′*(*t*). The correlation coefficients between 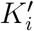 and *K*_*i*_ and between *x*(*t*) and *x′*(*t*) are close to 1.

The correlation coefficient between 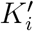 and *K*_*i*_ is further plotted versus the start time *t\** in Fig. 10(b). The correlation coefficient values remain above 0.90, though the one at *t\** = 45 minutes is slightly lower then others. This can be explained by the fact in Fig. 6(b) that the linear correlation between *x* and *x′* for *t\** = 45 minutes (i.e., the last three points) is relatively weaker, possibly due to higher noise in the cardiac scan. The corresponding *K*_*i*_ and 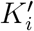 images for *t\** = 45 minutes are shown in Fig. 11. Overall, the two images still have very similar contrast appearance, indicating the scaling relation between *K*_*i*_ and 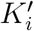.

**Figure 7.**
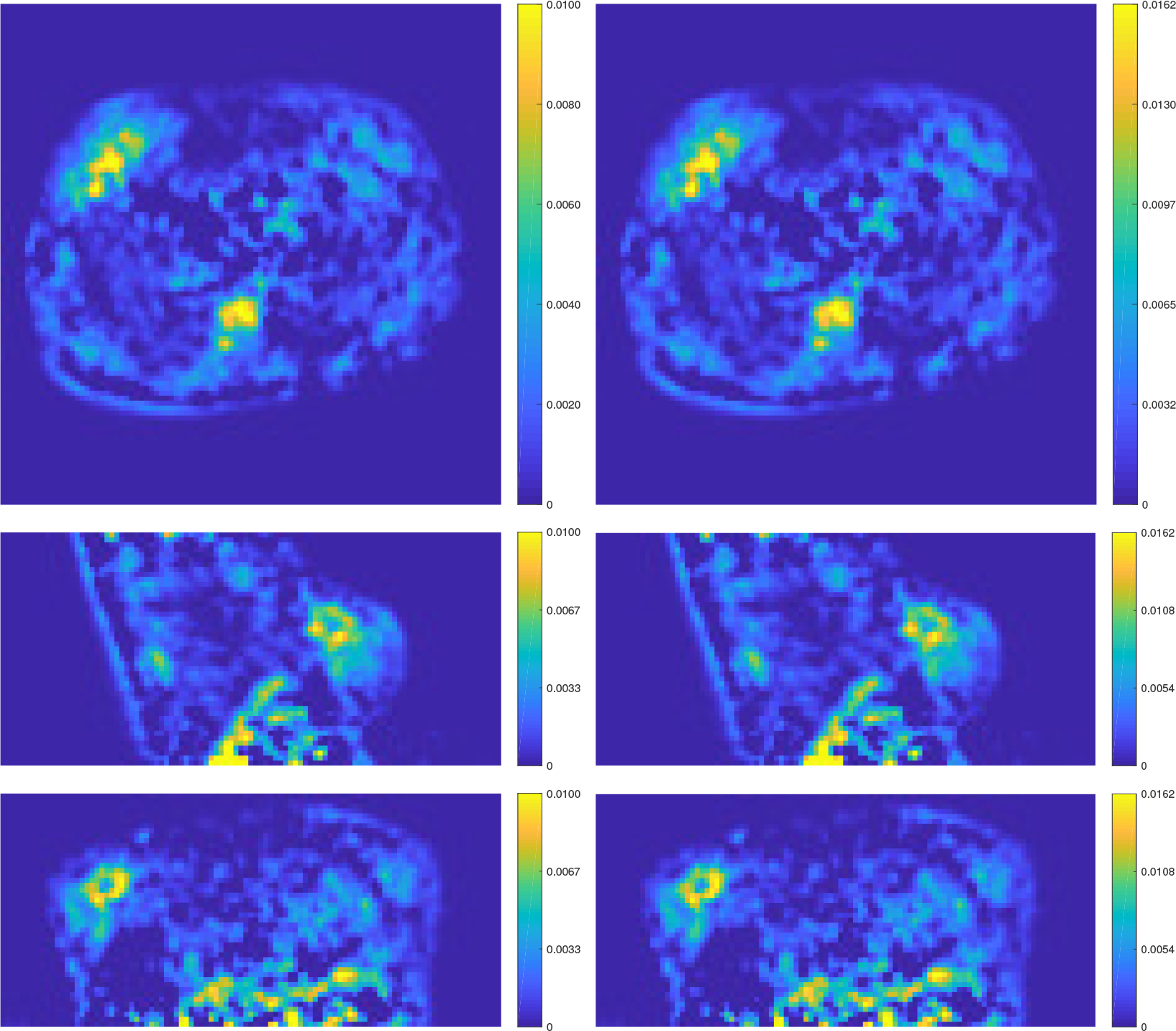
Comparison of parametric imaging using the standard Patlak plot and relative Patlak plot for the breast patient. Left: parametric image of the standard Patlak slope *K*_*i*_; Right: parametric image of the relative Patlak slope 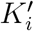. The start time *t\** = 30 minutes. From the top to the bottom are the views from the planes of transverse, sagittal and coronal, respectively.

**Figure 8.**
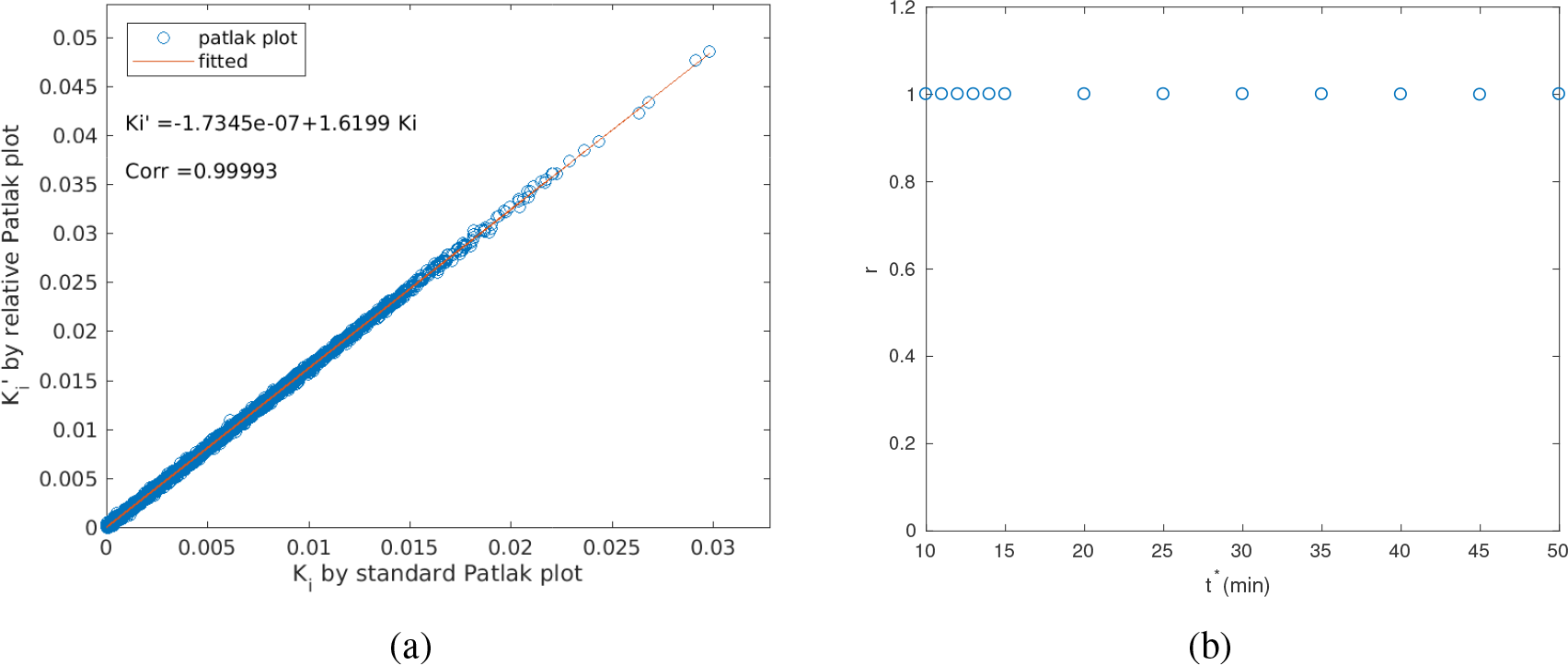
Results of the breast patient scan. (a) Relation between 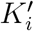, and *K*_*i*_ (*t\** = 30 minutes); (b) Correlation coefficient between 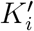, and *K*_*i*_, versus various start time *t\**.

**Figure 9.**
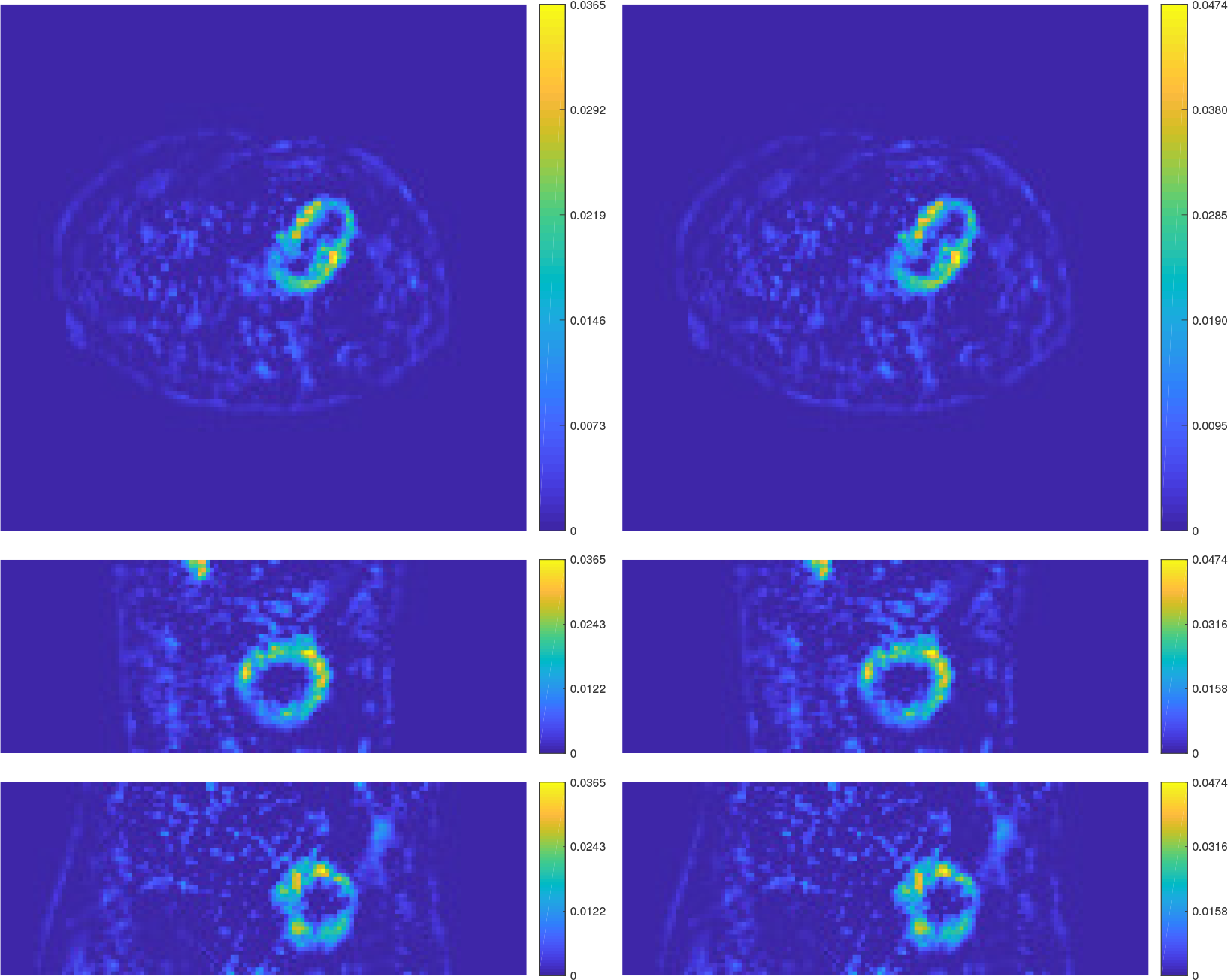
Comparison of parametric imaging using the standard Patlak plot and relative Patlak plot for the cardiac patient. Left: parametric image of *K*_*i*_; Right: parametric image of 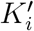. The start time *t\** = 30 minutes. From the top to the bottom are the views from the planes of transverse, sagittal and coronal, respectively.

## 5. Conclusion

We propose a new relative Patlak plot method for analyzing dynamic PET data. The new plot excludes the need for early-time input function and only requires late-time input function data, thus is easier to use than the standard Patlak method. Theoretical analysis, simulation results and real patient data all have demonstrated that parametric imaging by the relative Patlak plot determines the parametric image of the standard Patlak slope up to a global scaling factor. The new relative plot can replace the standard Patlak plot for certain applications where the determination of the global scaling factor is not necessary. Examples include, but are not limited to, lesion detection (e.g., [Li et al., 2009, Yang et al., 2016]) and metabolic tumor volume segmentation (e.g., [Visser et al., 2008]) using parametric map of tracer influx rate.

The investigation and development in this work are also timely because recently whole-body Patlak parametric image reconstruction has been adopted for commercial PET scanners [Hu et al., 2017]. The new relative Patlak plot can have a clear implication for practical use.

**Figure 10.**
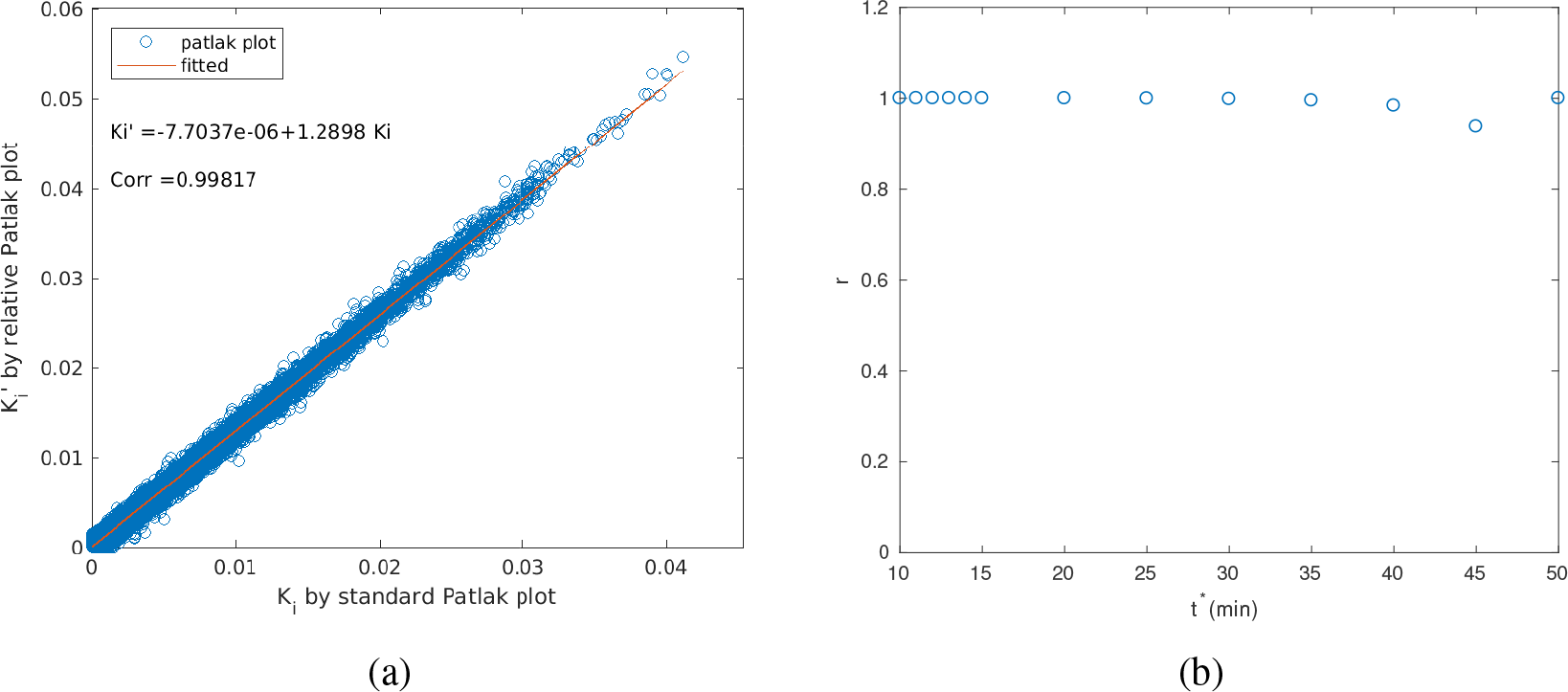
Results of the cardiac patient scan. (a) Relation between 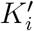 and *K*_*i*_ (*t\** minutes); (b) Correlation coefficient between 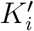 and *K*_*i*_ versus various *t\** values.

**Figure 11.**
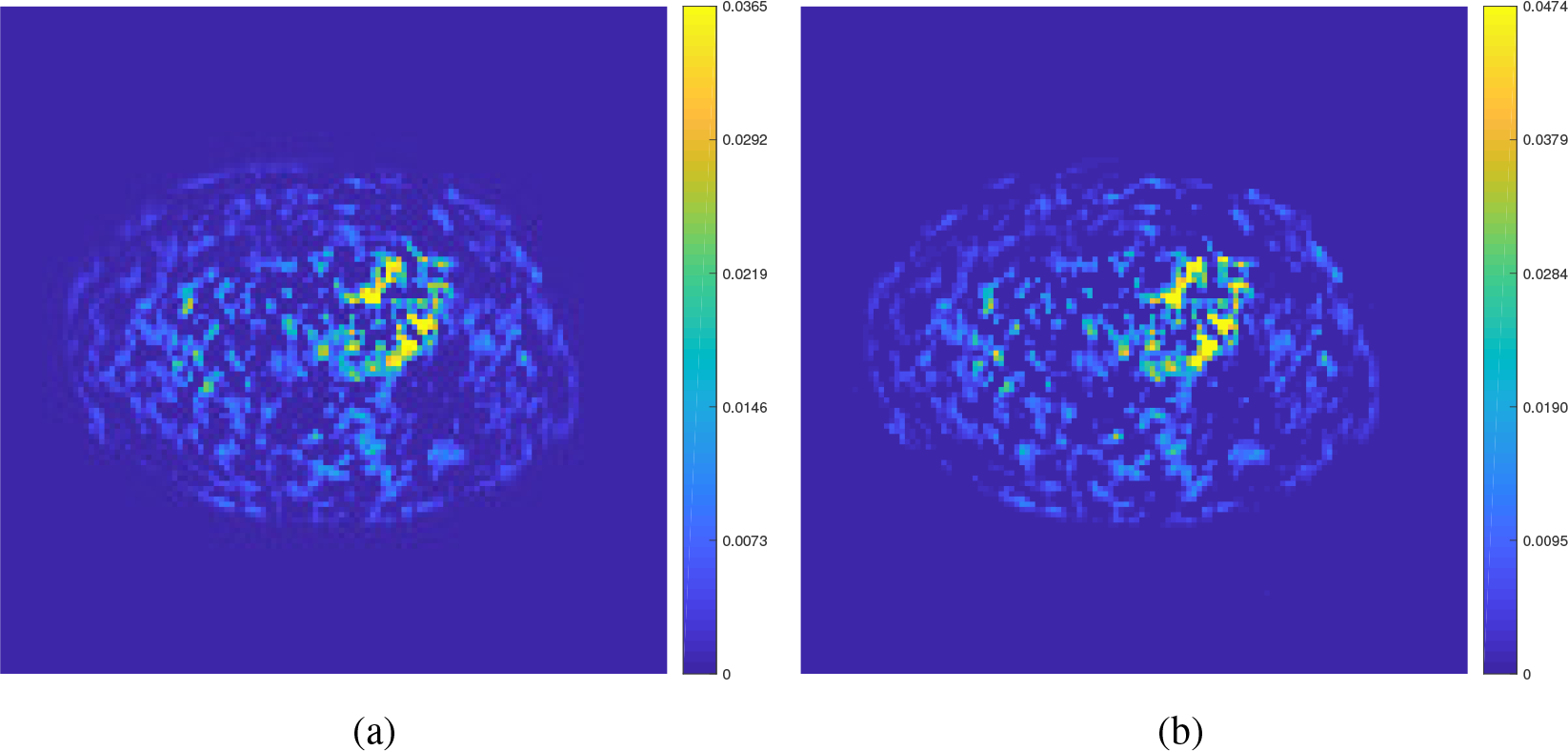
Parametric images of *K*_*i*_ and 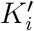 estimated with *t\** = 45 minutes for the cardiac patient data. (a) *K*_*i*_ by the standard Patlak plot; (b)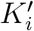 by the relative Patlak plot.

## Acknowledgments

The work is supported in part by NIH under no. R21 HL 131385 and by AHA under no. BGIA 25780046.

